# Injectable hydrogel embedded with mesenchymal stem cells repairs severe spinal cord injury

**DOI:** 10.1101/2022.07.01.498514

**Authors:** Xiangfeng Chen, Wujie Lu, Yanming Zuo, Jingjia Ye, Xiaodan Li, Zhonghan Wu, Shuang Jin, Wanxiong Cai, Zeinab Abdelrahman, Tianfang Zhang, Xiaosong Gu, Bin Yu, Zuobing Chen, Xuhua Wang

**Author notes:** These authors contributed equally (Xiangfeng Chen, Wujie Lu, Yanming Zuo, Jingjia Ye). Correspondence (X.W.), (Z.C.).

## Abstract

Mesenchymal stem cell (MSC) transplantation was suggested as a promising approach to treat spinal cord injury (SCI). However, the heterogeneity of MSC and the lack of appropriate delivery methods impede its clinical application. To tackle these challenges, we first generated human MSCs derived from a single cell with a great homogeneity of batch quality and then developed a biocompatible injectable hydrogel to embed these cells to treat severe SCI. In a clinically relevant rat severe SCI model, we showed that the injection of MSCs with injectable hydrogel into the lesion site promoted robust functional recovery, while the intrathecal delivery of MSCs only resulted in limited therapeutic effects. Mechanistically, the hydrogel protected MSCs from the damage of harmful neuroinflammatory microenvironment in the spinal cord lesion. The hydrogel with the survived MSCs ameliorates the neuroinflammatory microenvironment of spinal cord lesion, preventing cavity formation and leads to the remnant of spared axons/tissues, which results in a better prognosis in the end.

## Introduction

Severe traumatic spinal cord injury (SCI) leads to tragic neurological dysfunction, a decrease of life quality, and a serious financial burden^1^. However, there is no efficient treatment for these neurological deficits nowadays^2^. Following SCI, the penetration of immune cells into the injury site causes a dysregulated inflammatory microenvironment that induces neuronal/glial apoptosis and inhibits nerve regrowth^3, 4^. As the primary mechanical injury is unpredictable, the following secondary injury due to neuroinflammation is the main target for clinical treatment^5^.

Recently, mesenchymal stem cells (MSCs) have been proven effective in treating spinal cord injury because of their ease of preparation and outstanding properties of anti-inflammatory and minimal immunoreactivity^6^. However, two major obstacles prevent MSCs from clinical application. First, the heterogeneity of patch quality of the produced MSCs is one of the major obstacles to their clinical application. MSCs could originate from numerous tissues, including embryonic stem cells^7^, bone marrow^8^, umbilical cord^9, 10^, adipose^11^ and amniotic fetal^12^, which results in a huge heterogeneity during MSC production, thus developing a method to produce MSCs with controllable patch quality is crucial and challenging. Second, an appropriate delivery method is highly demanded to support MSCs survival in the neuroinflammatory microenvironment of a spinal cord lesion, which significantly limits their therapeutic effects^13^. To overcome this limitation, numerous biomaterials, including synthetic biodegradable hydrogels (such as polylactic acid (PLA), polyglycol acid (PGA), and polyethylene glycol (PEG), natural scaffolds (such as fibrin, collagen and hyaluronic acid) have been suggested to encapsulate MSCs for SCI treatment^14, 15^. However, the synthetic hydrogels are still suffering from poor biocompatibility for the loaded cells, and the injectable natural scaffolds which can reduce injury during operation might be more suitable for SCI treatment^16^.

To address the above challenges, we proposed an injectable hydrogel embedded with human embryonic stem cell derived MSCs (hESC-MSCs) to repair severe SCI by preventing cavity formation and regulating the neuroinflammatory microenvironment of spinal cord lesions. Because the produced hESC-MSCs in our study were derived from one single cell, they are with remarkable homogeneity of batch quality. The proposed injectable scaffolds were based on natural materials with great biocompatibility. In addition, the injectable hydrogel can fill the irregularly shaped injury cavity and support cell survival. More importantly, the hydrogel could easily load various therapeutic agents (e.g., anti-neuroinflammatory drugs and cytokines) to ameliorate the microenvironment and facilitate spinal cord repair. Taken together, we provide a method to deliver MSCs with an excellent homogeneity for translational SCI study.

## Results

### Characterization of standardized MSCs derived from human embryonic stem cells

To eliminate the heterogeneity of batch quality of produced MSCs, we produced MSCs from human embryonic stem cells as the flow chart (Fig. 1A) with a method adapted from a previous study^17^. The produced MSCs were then analyzed by flow cytometry with the MSC specific markers of CD73, CD105, and CD90, as well as control markers of CD14, CD34, CD45, CD79a and HLA-DR. As shown in Fig. 1B, the markers specific for MSCs exhibited more than 99% positive, while the expressions of the control markers, CD14, CD34, CD45, CD79a and HLA-DR, were extremely low. Moreover, the morphology of the produced MSCs with different times of passage was almost the same (Fig. 1C), indicating the high homogeneity of the produced MSCs. Additionally, we further evaluated the expression of the ESC-specific transcription factors (TFs) SOX2, OCT4 and NANO by qRT-PCR, confirming that the produced MSCs showed less expression of the three TFs compared to umbilical cord-derived mesenchymal stem cells (ESC-MSCs) or human embryonic stem cells (ESCs) control (Fig.1D). The results suggest that the produced hESC-MSCs consistently exhibit typical MSC characteristics at both the transcriptional and morphological levels.

**Figure 1.**
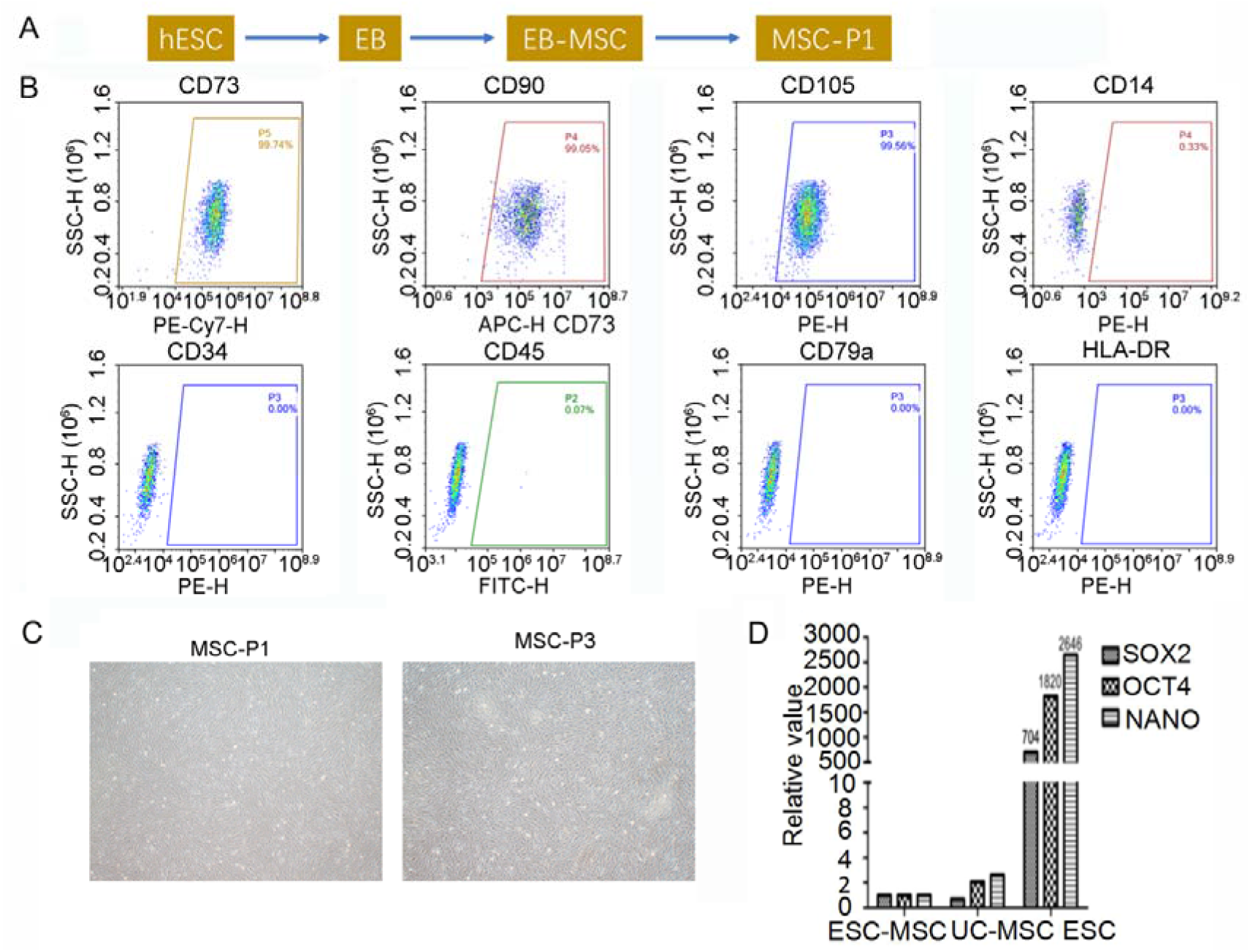
Production and characterization of MSCs from human embryonic stem cells (hESC-MSC). (A) Schematic of the process to obtain MSCs from human embryonic stem cells (hESC-MSC). (B) Representative bivariate plots of MSCs determined by flow cytometry analysis (FACSs). (C) Representative cell pictures of MSCs from different passages. (D) RT-qPCR analysis revealed the mRNA expression levels of stem cell specific transcription factor gene SOX2, OCT4 and NANO. ESC-MSC, embryonic stem cell derived mesenchymal stem cell. UC-MSC, umbilical cord derived mesenchymal stem cells. ESC, embryonic stem cell.

### Intrathecal injection of hESC-MSCs improved limited functional recovery after severe SCI

As the intrathecal injection is a quite acceptable delivery method in clinical and some previous works have shown that intrathecal injection of MSCs could inhibit neuroinflammation and promote neuron/tissue survival after SCI, we attempted to evaluate whether intrathecal injection of the produced MSCs could benefit the functional recovery of rats with severe SCI.

We first established a clinically relevant model of severe SCI induced by a contusion at the level of the tenth thoracic (T10) vertebra as in our previous work^18^. Then intrathecal injections of 20 μl PBS or PBS with MSCs (total ~ 1× 10^6^ cells) were administered to rats at the same day after SCI (Fig. 2A). To visualize the propriospinal axons descending from rostral segments to caudal segments, we injected AAV2/9-hsyn-mCherry into the upper thoracic spinal cord (T8) 2 weeks before perfusion (Fig. 2A).

**Figure 2.**
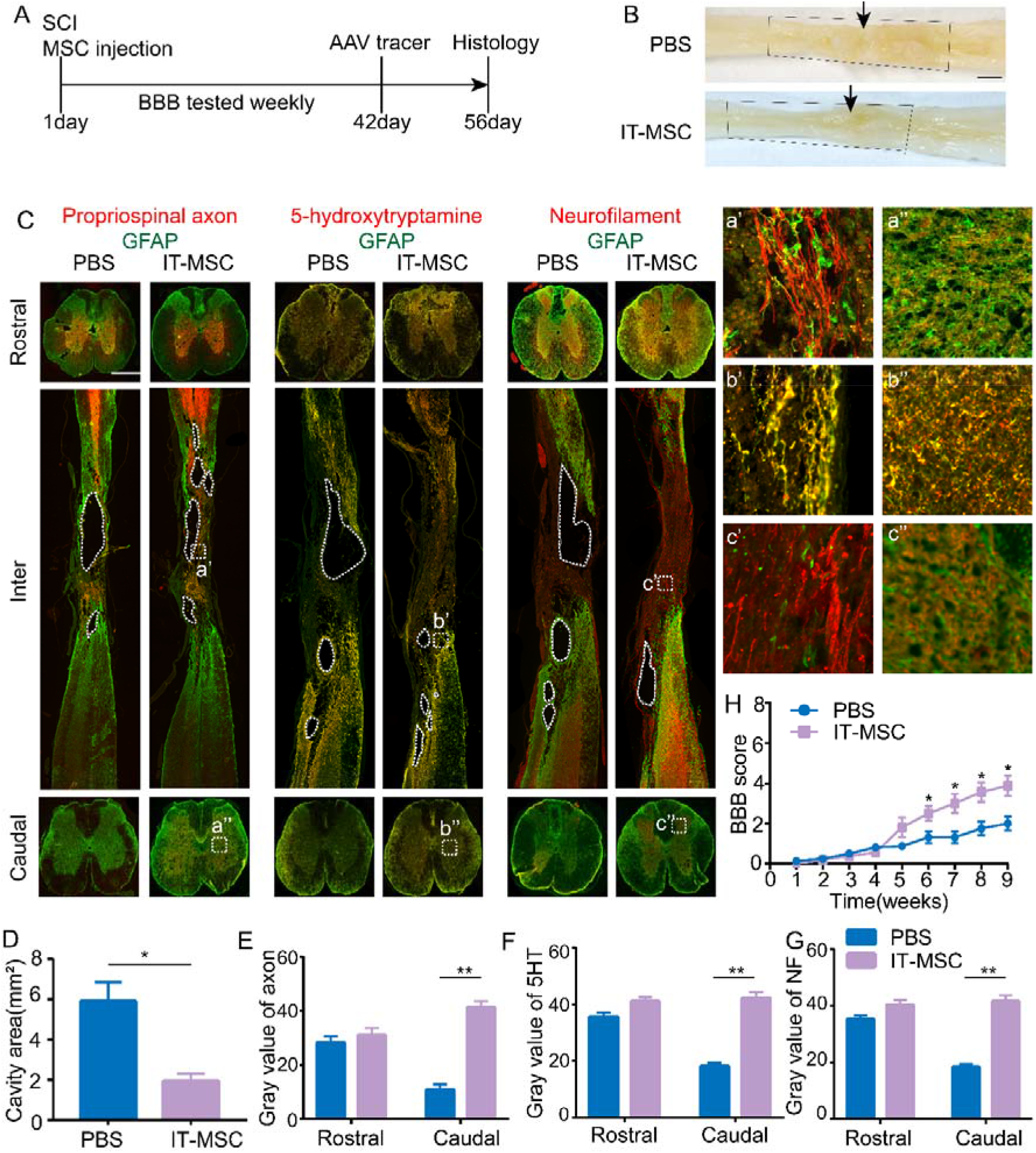
MSCs treatment through intrathecal injection reduced cavity areas and resulted in limited function recovery in SCI rats. (A) Schematic diagram of the experimental design. PBS or MSCs were injected intrathecally within 4 h after SCI. The axon tracer was injected at 6 weeks after SCI, and the animals were finally sacrificed at 8 weeks after injury. (B) Whole spinal cord images at 8 weeks after SCI. Black arrows indicate the injury site. The dotted boxes indicate the expected normal shapes of the injured spinal cord. (C) Representative images of immunofluorescent staining for GFAP (green), 5-HT, propriospinal axons, and NF (red) under different injection conditions after 8 weeks. Coronal sections rostral (top row) and caudal (bottom row) to the injury site, showing the axon density after injection. Sagittal sections (middle row) showing the cavity and spared tissue in the injury site. Dotted lines indicate the cavity. a’, b’, and c’ show the details of scar formation at the injury site. a”, b”, and c” show enlarged images of the axon density on the caudal side after injections. (D) Quantification of the cavity areas on the sagittal sections at the injured sites,.Values were expressed as mean ± SD, n = 5. (E-G) Quantification of propriospinal axon (E), 5-HT positive axon (F), and NF density (G) on the rostral and caudal sides by the gray value; all values were expressed as the mean ± SD, *n* = 5. (H) Comparison of locomotion recovery among the different groups (PBS, IT-MSC) measured using the BBB scale each week in an open field, the values were expressed as mean ± SEM, n = 10. (D–H) Statistical analysis was performed using unpaired, two-tailed Student’s t test \**P* < 0.05 and \*\**P* < 0.01.

At the 8^th^ week after injury, we found that the spinal cord of both groups showed a remarkable shape deformation (Fig. 2B), and in both groups, cavities without any tissue matrix were presented at the injury site (Fig. 2C, inter, dotted line areas). However, the PBS group showed large cavities extending rostrally and caudally more than 4 mm away from the epicentre of the injury site, while the rat with MSC intrathecal injection (IT-MSC) group showed smaller cavities (Fig. 2B, C, D). The cavity area in the IT-MSC group was nearly one third of that in the PBS-injected control group (Fig. 2D).

In the spinal cord sections, propriospinal axons, descending serotonergic axons, and ubiquitous nerve fibres (Figure 2C) were visualized by immunohistochemistry with an anti-red fluorescent protein (RFP), anti-5-hydroxytryptamine (5-HT), and anti-neurofilament (NF) antibodies, respectively (Fig. 2C). We found that injured axons stopped before the host/cavity borders with intense glial fibrillary acidic protein (GFAP)+ scarring, and no axons extended into the cavity in any of the examined subjects (n= 10) (Fig. 2C, inter). Coronal sections were prepared and stained with same antibodies to assess the quantity of spared axon across the lesion. As expected, no significant difference was observed in any axon in coronal sections of spinal segments above the injury site between the two groups (Fig. 2C, rostral). However, there were more propriospinal, 5-HT, and NF^+^ axons at the injury sites of IT-MSC treated rats compared to that of PBS-treated rats (Fig. 2C, a’, b’, and c’). Correspondingly, coronal spinal imaging and quantification of the axon density below the injury site showed that there were more propriospinal, 5-HT, and NF^+^ axons caudal to the injury site in the IT-MSC treated rats (Fig. 2C, caudal and 2E–2G). These beneficial treatment effects possibly contribute to the down-regulation of the inflammation during SCI induced by the injected MSCs, as shown by the decreased expression of microglia (CD68^+^) and glial scars (GFAP^+^) (Fig. S1A, S1B and S1E). Moreover, fibronectin (FN) and NeuN staining of coronal spinal sections at the injured sites indicated that the cavities formed in control animals were largely filled by FN^+^ matrix and more NeuN^+^ neuron were observed in IT-MSC treated rats (Figure S1A, S1C and S1D). These results indicate that IT-MSC could promote spared neurons survival and provide beneficial FN^+^ matrix at the injured sites.

Unfortunately, the hindlimb locomotor functional recovery of the IT-MSC treated rats was limited, as indicted by the Basso, Beattie, and Bresnahan (BBB) scores of the rats (Fig. 2H). The BBB scores of IT-MSC treated rats were only elevated by 1-2 compared to the PBS group (Fig. 2B), indicating that intrathecal injection of MSCs only leads to minimal therapeutic effects on spinal cord injury. This observation was associated with the limited number of spared axons rescued by MSC treatment through this method (Fig. 2C) and the relatively short survival time (2 weeks) of MSCs after intrathecal injection (Fig. S2F and S2G).

### Design and Fabrication of Self-Assembling Hydrogel Depot

Since the intrathecal injection of MSCs only achieved minimal therapeutic effects on SCI rats, we next sought to examine whether delivery of MSCs locally to the SCI lesion site could result in more substantial therapeutic effects. However, the injected MSCs might not survive for a long time in the inflammatory environment of SCI lesion^19^. Previous studies have shown that injection of hydrogel into the injured site of the spinal cord could successfully fill the irregularly shaped SCI lesions, dramatically reduce the shape deformation and prolong the survival of MSCs^20^. For this purpose, the injectable gel should be biocompatible and facilitate the incorporation of anti-inflammation treatment. To this end, we designed an injectable, natural biological macromolecule-based, self-assembling hydrogel depot with the capacity to be loaded with different therapeutic agents. Because gelatin has excellent biocompatibility, we used this material as the backbone to generate the hydrogel depot (Fig. 3A). First, the commercial gelatin was sulfhydrylated, utilizing its free hydroxyl and amino groups to get gelatin-SH (Fig. S2A and S2B). As for the linker, we chose 2arm-PEG-maleimide (2a-PEG-MAL, M.W. 20,000), which could link gelatin-SH to create a 3D polymer network, and mimic the native extracellular matrix (ECM) through thiol-maleimide crosslinking (Fig. 3A). Through the Michael addition reaction between the thiol group and maleimide, a self-assembling hydrogel depot was formed in less than 5 s at a concentration of 5% 2a-PEG-MAL and 8% gelatin-SH (defined as 5% Gel in brief), and gelation was much slower at a lower concentration of 2a-PEG-MAL (Fig. 3E). To figure out whether the resulting hydrogel could support MSCs survival, we loaded MSCs into hydrogels with various concentrations of cross-linker followed by an MTT assay for cell proliferation detection and fluorescence imaging of cells with death cell staining (Fig. 3J and 3K). The results showed, that the 4% and 5% Gel are with elastic moduli of ~ 220-460 kPa, which resembles normal spinal cord tissue (200-600kPa)^21^, but induces no significant cell death (Fig. 3K). Comprehensively considering their gelatinizing performance, we chose 5% Gel for the following in vivo assays. Swelling studies of the 5% Gel showed a low equilibrium swelling ratio of 15% after immersion in PBS for the first day and slightly increased to a 25% plateau in the following 5 days (Fig. 3F). This swelling ratio is acceptable, and this material is unlikely to initiate second extra damage because it is similar to the natural materials such as HA and fibrin, which have been suggested as suitable materials for transplantation^22^. We further investigated the degeneration profile of the 5% Gel by measuring the mass loss of lyophilized samples for 1 month (Fig. 3G). The results showed that the 5% Gel exhibited relatively slow degradation during the first 3 days and dramatically reduced to 35% at 7 days. After 2 weeks, the material degraded about 90%, and the degradation level reached a plateau during the next 2 weeks, indicating that the hydrogel depot can support cell migration for about 2 weeks to 1 month after injection and be quickly replaced by real ECM to better support cell survival and tissue repairing after SCI.

**Figure 3.**
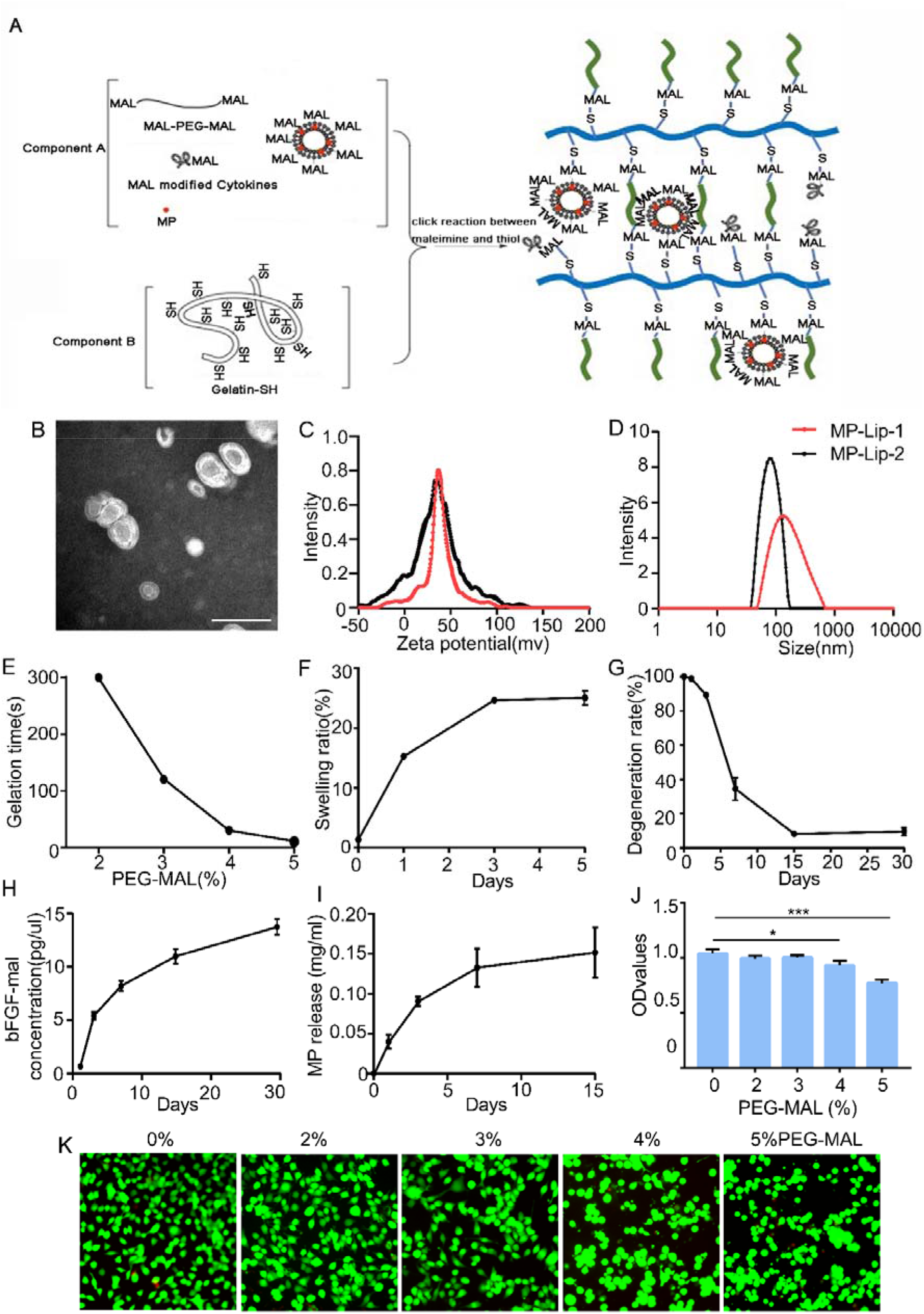
Fabrication and identification of self-assembling hydrogel depot. (A) Schematic of the synthesis of the gelatin-based injectable hydrogel and visualization of the gelation process by the mixture of solution A and solution B. (B) SEM of liposomes loaded with MP. (C) Zeta potential distribution of liposomes loaded with MP. (D) Size distribution of liposomes loaded with MP. (E) Gelation time of 8% Gelatin-SH with different concentrations of MAL-PEG-MAL. (F) Swelling ratio of the gelatin-based hydrogel in PBS at 37 °C from 0–5 days, data are presented as mean ± SD (n=3). (G) Degeneration rate of the gelatin-based hydrogel in PBS at 37 °C for 30 days, data are presented as mean ± SD (n=3). (H) Release of bFGF from the hydrogel in PBS at 37 °C for 30 days, data are presented as mean ± SD (n=3). (I) Release of MP from the hydrogel in PBS at 37 °C from 0–15 days, data are presented as mean ± SD (n=3). (J) Cytotoxicity of the hydrogel tested by MSC survival using MTT assay. Data are presented as mean ± SD (n=3), and statistical analysis was performed using unpaired, two-tailed Student’s t test \**P* < 0.05 and \*\*\**P* < 0.001. (K) Cytotoxicity of the hydrogel tested by MSC survival using cell death staining.

In addition, considering the scarcity of nutrient supply and hyperactive neuroinflammatory environment in the SCI lesion, we co-loaded an anti-inflammatory drug and a group of growth factors (GFs) into the hydrogel depot. As we know, the hydrogel depot is difficult to load hydrophobic drugs, while many clinically available therapeutic drugs such as prednisone and methylprednisolone (MP) are hydrophobic. To overcome this difficulty, we encapsulated the hydrophobic anti-inflammatory drug MP in MAL-modified liposomes to get hydrophobic drug-loaded liposome nanoparticles (MP-lip-MAL) (Fig. S2C), such that MP could be released from the hydrogel depot in a sustained manner (Fig. 3I). Scanning electron microscopy (SEM) was used to visualize the resulting MP-lip-MAL, indicating that the nanoparticles were spherical with an average size of about 100 nm (Fig. 3B). The zeta potential of the MP-lip-MAL was about 20 mV (Fig. 3C), and the particle size of the MP-lip-MAL was about 100 nm or 200 nm depending on the MP-loading rates measured by dynamic light scattering (Fig. 3D) and the 100 nm MP-lip-MAL liposomes were chosen for the following *in vivo* assay because they are relatively stable than large liposomes. Importantly, when the MP-lip-MAL were conjugated to the 5% Gel, MP was released from the hydrogel for more than 15 days in a sustained manner (Fig. 3I), which could continuously benefit the anti-inflammation therapy without bringing side effects caused by systemic administration. As for the cytokines, we chose human brain-derived neurotrophic factor (BDNF) to support neuron survival and recombinant human vascular endothelial GF (VEGF) and recombinant human fibroblast growth factor (bFGF) to promote angiogenesis and conducted maleimide-modification as previously reported^23^. Moreover, when the maleimide-modified bFGF (bFGF-MAL) was loaded into the 5% Gel, the conjugated bFGFs were continually released for more than 1 month (Fig. 3H), which could provide sufficient support for tissue or cell survival as well as the survival of loaded MSCs in the chronic phase of SCI. Based on the above results, the 5% Gel is permissive for MSCs loading and shows various beneficial characteristics for injection into the injury sites of the spinal cord.

### Hydrogel depot with MSCs and synergistically released GFs, and anti-inflammatory drug prevents cystic cavity formation and facilitates nerve regrowth

Previous studies have shown that neurotrophic factors and anti-inflammatory drugs could support MSCs survival and promote nerve regrowth^24^. Thus, we speculated that MSCs loaded into the 5% Gel with synergistic released GFs and an anti-inflammatory drug might prolong MSCs survival, promote spared neuron/tissue repair and prevent cystic cavity formation. To test this hypothesis, we first evaluated the survival of MSCs injected into the injury sites of the SCI rats 1 week after the injury, and SCI animals injected with PBS served as control. As shown in Fig. S3A and S3B, although the survived MSCs reduced with the extension of time, the injected MSCs could survive for more than 8 weeks, which was significantly longer than that of the intrathecally injected MSCs (2-3 weeks, Fig. S1F and S1G). Next, MP-lip-MAL and MAL-GFs loaded with 5% Gels with or without MSCs (Gel or Gel +MSCs) were injected into the injury site as described in the method section at 7 days post-injury (Fig. 4A). Notably, compared with the remarkable shape deformation in the PBS or IT-MSC treated animals (Fig. 2B and 4B), Gel or Gel+MSCs injection remarkably reduced shape deformation (Figure 4B). Moreover, the cystic cavities induced by SCI were much smaller in the rats with Gel or Gel+MSCs treatment, especially for the Gel+MSCs rats whose cystic cavities almost disappeared (Fig. 4C, Inter). In addition, after Gel or Gel+MSCs treatment, a few red mCherry-labeled propriospinal and serotonergic (5-HT) positive axons grew and extended into the lesion site, as visualized by microscopy in all examined subjects (n = 8) (Fig. 4C left and middle, a’, b’). More strikingly, large numbers of ubiquitous NF^+^ axonal fibres regrew into the GFAP-negative matrix and extended into the whole injury site in Gel or Gel+MSCs treated rats, especially for Gel+MSCs (Fig. 4C right, c’). Correspondingly, the density of red mCherry-labeled propriospinal and serotonergic (5-HT) positive axons, as well as NF^+^ axon, below the injury site in the gel treated animals was significantly higher than that of the PBS control group. In contrast, the Gel+MSCs treated rats exhibited even more axons than the gel treated rats (Fig. 4C caudal, a’’, b’’, c’’, 4D–4F), indicating the beneficial therapeutic effects of both Gel and MSCs in SCI treatment.

**Figure 4.**
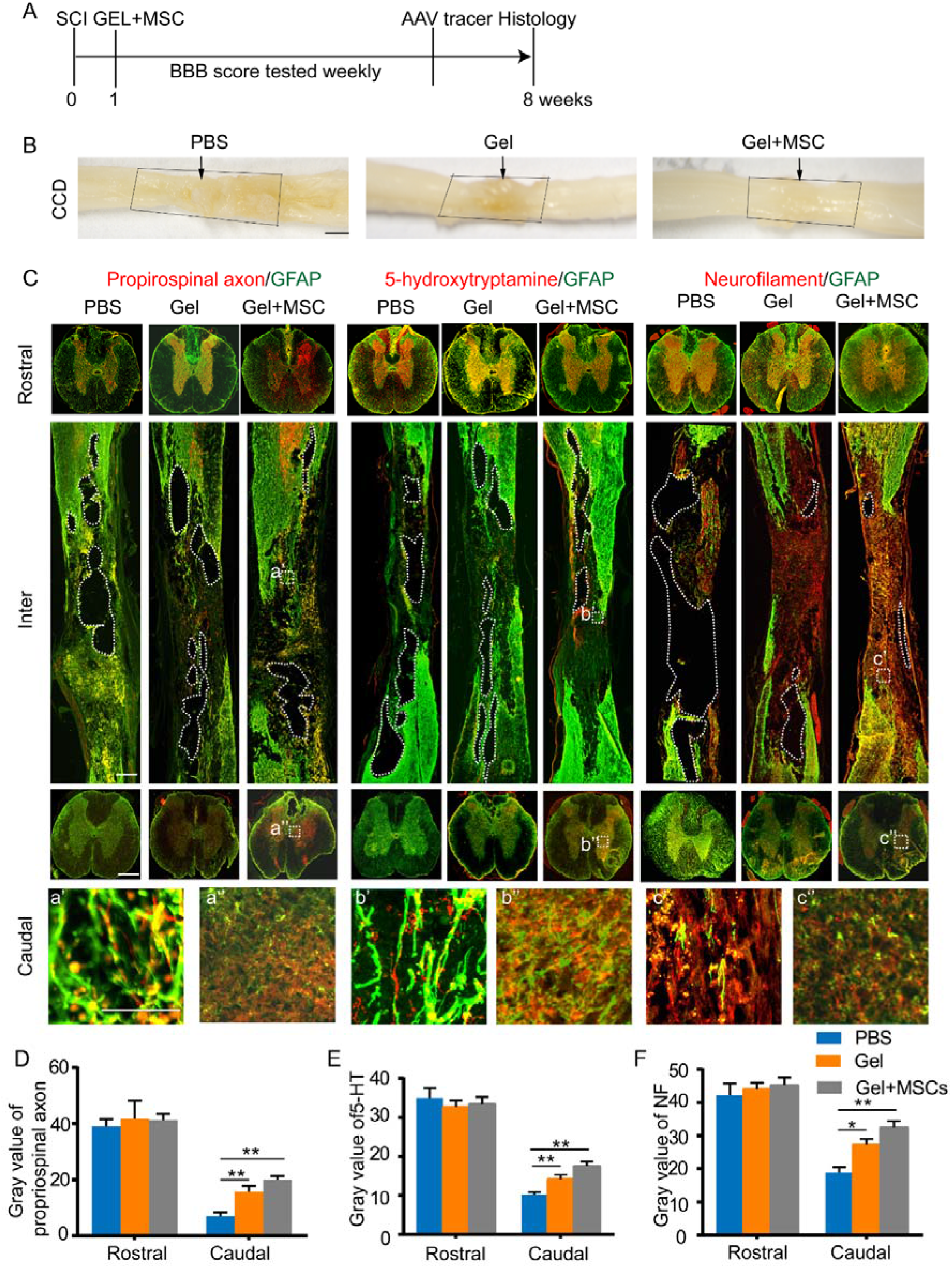
Mercapto gelatin-based hydrogel depot with MSCs and synergistically released GFs and anti-inflammatory drug prevents cystic cavity formation and facilitates nerve regrowth. (A) Schematic diagram of the experimental design. PBS, Gel or Gel+MSCs were injected into the injured site at 1 week after SCI. The axon tracer was injected at 6 weeks after SCI, and the animals were finally sacrificed at 8 weeks after injury. (B) Whole spinal cord images at 8 weeks after SCI. Black arrows indicate the injury site. The dotted boxes indicate the expected normal shapes of the injured spinal cord. (C) Representative images of immunofluorescent staining for GFAP (green), 5-HT, propriospinal axons, and NF (red) under different injection conditions after 8 weeks. Coronal sections rostral (top row) and caudal (bottom row) to the injury site, showing the axon density after injection. Sagittal sections (middle row) showing the cavity and spared tissue in the injury site. Dotted lines indicate the cavity. a’, b’, and c’ show the details of scar formation at the injured site. a”, b”, and c” show details of the axon density on the caudal side after injections. (D-F) Quantification of propriospinal axon (E), 5-HT positive axon (F), and NF density (G) on the rostral and caudal sides by the gray value; data are presented as mean ± SD (n=8). Statistical analysis was performed using unpaired, two-tailed Student’s t test \**P* < 0.05 and \*\**P* < 0.01.

### Gel or Gel+MSCs treatment promotes functional recovery through regulating neuroinflammation

We next examined whether the observed beneficial therapeutic effects could lead to functional recovery. After analyzing the recorded behavior performances of the rats in a double-blind manner, we found that the Gel and Gel+MSCs treatment significantly improved angular mobility during the first 2–3 weeks after treatment. Through the BBB scoring, we found that Gel and Gel-MCSs groups improved the motor ability of animal hind limbs, from 2-3 points to 6-8 points. After analyzing the gait of animal hind limbs, we found that the total stride of hind limbs in Gel and Gel+MSCs treatment groups increased significantly, and there were more steps and slight joint activity compared with the PBS group. However, compared with the intact group, these activities are still incomplete, indicating that the recovery of these rats is not ideal. In the analysis of rat joint angle, it was observed that the animals in Gel and Gel+MSCs groups had a slight bend in joint angle, indicating that the hind limbs of the animals had a certain motor ability, but these movements were far from those in the normal group.

Similarly, in Gel and Gel+MSCs treated rats, stride improvement and weight support were observed. More importantly, the Gel+MSCs group showed a significant difference compared with the gel group, suggesting that the MSCs in the gel supported SCI repair and enhanced function improvement (Fig. 5E). In addition, to figure out the mechanism by which the hydrogel and MCSs repaired spinal cord injury, we investigated the gene expression in injured spinal cord tissues by real-time PCR. We analyzed several genes, IL-6, TNF-a, IL-4 and IL13, related to the inflammatory response. The results showed that Gel and Gel+MSCs treatment protected spared axons from secondary injury by inhibiting the expression of pro-inflammatory genes IL-6, and TNF-a and promoting the expression of anti-inflammatory genes IL-4 and IL13 (Fig.5F). Through this mechanism, the spared neural circuits were protected and reorganized, such that the motor ability of the hindlimb in paralyzed rats was partially restored.

**Figure 5.**
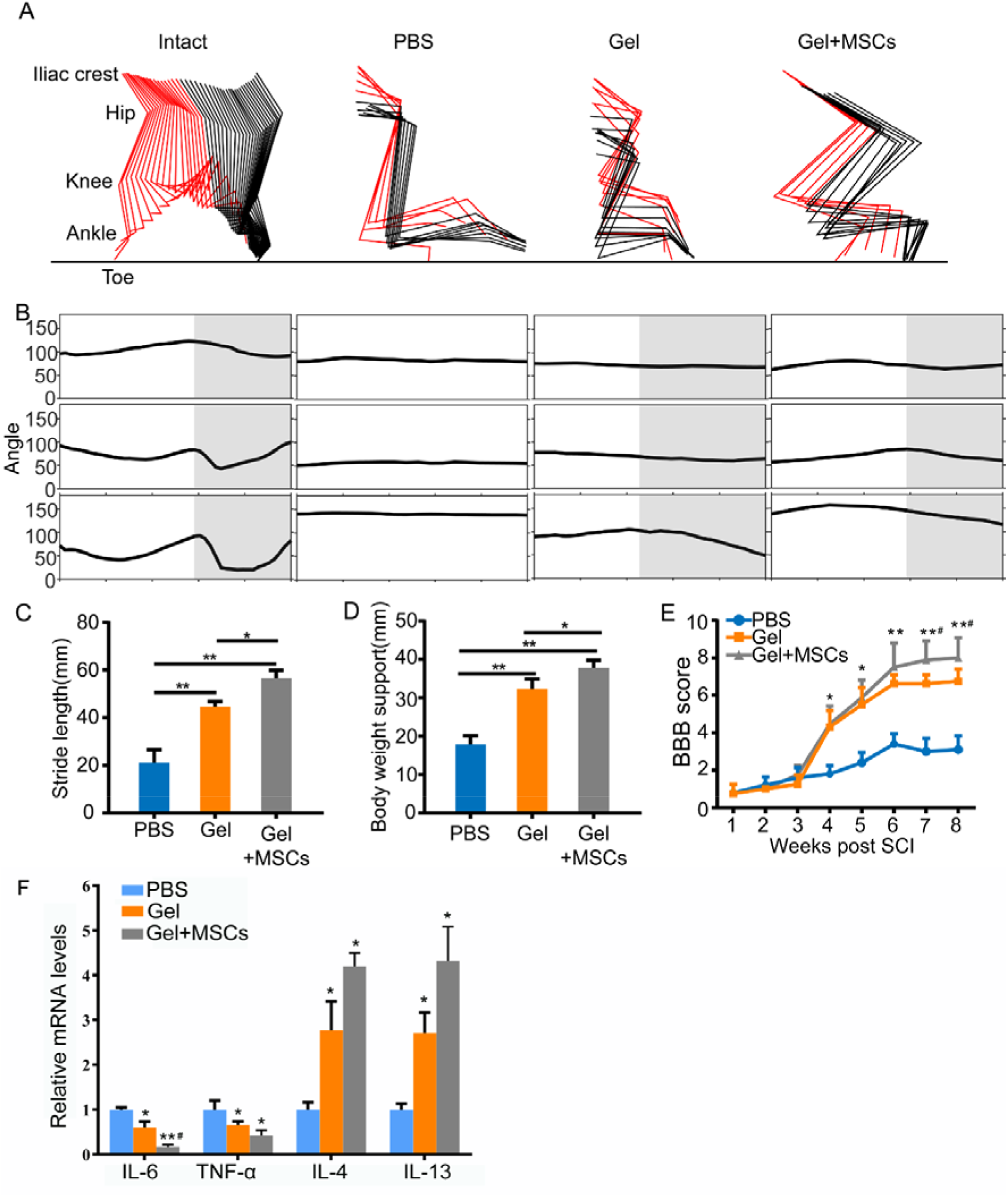
Gel or Gel+MSCs treatment mediates functional recovery through regulating neuroinflammation. (A-B) Rat hindlimb movement (A) and Angle movement curve of various groups showed various color-coded stick drawing (intact, PBS, Gel and Gel+MSCs). (C-D) Quantification of stride length support (C) and body weight support (D) at 8 weeks. Data are presented as mean ± SEM (n=3). Statistical analysis was performed using unpaired, two-tailed Student’s t test. \**P* < 0.05 and \*\**P* < 0.01. (E) Comparison of locomotion recovery among the different groups (PBS, Gel and Gel+MSCs) measured using the BBB scale each week in an open field, data are presented as mean ± SEM (n=10). Statistical analysis was performed using unpaired, two-tailed Student’s t test *\*P* < 0.05 and \*\**P* < 0.01 was compared to the PBS group. #*P*<0.05 was compared to the Gel group. (F) The expression levels of IL-6, TNF-a, IL-4 andIL13 at the injured spinal cord detected by real-time PCR. Data are presented as mean ± SD (n=3). Statistical analysis was performed using unpaired, two-tailed Student’s t test \**P* < 0.05 and \*\**P* < 0.01 was compared to the PBS group. #*P*<0.05 was compared to the Gel group.

## Discussion

Spinal cord injury is a pernicious disease affecting numerous families worldwide, but without effective treatment. One of the most challenging problems is to control the immune disorder of the injured site and fill the injured cavity. It is worth noting that the injectable scaffold combined with hESC-MSCs and MP-liposome described here supports the repair of severe SCI and results in robust functional recovery of hindlimb locomotion.

Nowadays, mesenchymal stem cells (MSCs) have attracted particular attention for their weak immunogenicity and significant anti-inflammatory effect in the spinal cord lesion^25^. MSCs can be induced from different animals and tissues, but various tissue-derived MSCs tend to have different characteristics, resulting in uncontrollable effects in the treatment of spinal cord injury^26^. Thus, the inconformity of MSCs seriously impedes their clinical application. In this study, we generated standardized MSCs cells derived from human embryonic stem cells, which can be passed for more than 20 times and have excellent stability and anti-inflammatory activity, facilitating translation of these experimental results into human clinical study.

Intrathecal injection of MSCs has been widely used in the treatment of spinal cord injury due to its advantages of convenient operation and less additional injury^27, 28^. However, its function in promoting spinal cord injury repair is controversial^29^. In this study, we compared and analyzed the treatment effect between intrathecal injection of MSCs and in-situ injection of MSCs, and found that intrathecal injection of MSCs could only reduce spinal cord cavity, promote injury repair and improve the motor ability to a deficient degree. The treatment efficacy of intrathecal injection of MSCs was significantly weaker than that of in-situ injection of MSCs, indicating that intrathecal injection of MSCs is suitable for mild spinal cord injury rather than severe spinal cord injury.

Compared with implantable scaffold materials, injectable gel materials have less damage to spinal cord tissues and do not require surgical implantation, thus with better operability and a wider application range^30^. However, many chemosynthetic injectable materials have the inherent defect of poor biocompatibility, making it difficult to be compatible with cell therapy or acceptable in clinical application^18^. Here we used gelatin-based and liposome-based drug delivery systems, resulting in injectable gel with good cell affinity. In previous reports, injection of different self-assembling peptides or hyaluronan-based hydrogels reduced cavity volume or promoted tissue preservation following clip compression injury, but sizable cystic cavities remained^30, 31^. In contrast to these studies, our gelatin-based hydrogel almost eliminated cavity spaces. In addition, cystic cavities in most human SCIs are highly irregular and unpredictable, with remaining more or less uninjured axons. Many previous studies tested injectable biomaterials in surgically created injury models that have a designed dimension of the injury site, making it difficult to evaluate their potential in the clinical application^32, 33^. Using a clinically relevant contusion injury model in rats, where fluid-filled cystic cavities progressively develop in a manner closely resembling what occurs after human SCI^34^, our study demonstrated that injection of the gelatin-based hydrogel could induce near-complete bridging of posttraumatic cavities.

However, we still need further monkey experiments to evaluate the therapeutic effect and optimize the dose before human clinical studies. Moreover, it has been proved that transcranial magnetic and functional electrical stimulation improve function in SCI patients^35^. So, we believe that combining proper training in the following clinical study with MSCs treatment could further improve the treatment effect.

## Experimental Section

### Animals

Female Sprague–Dawley rats (200-250 g, grade II, certificate no. SCXK2008-0033) were obtained from the Zhejiang Academy of Medical Sciences. All experiments were approved by the Zhejiang University School of Medicine Animal Experimentation Committee (approval ID: ZJU20210110) and were in complete compliance with the National Institutes of Health Guide for the Care and Use of Laboratory Animals. The spinal contusive injury was inflicted using an Infinite Vertical impactor (68099, RWD Life Science, China).

### Preparation of MSC

Human ESC (embryonic stem cell) derived MSCs were generated by Ysbiotech, Hangzhou, China (YS™ hESC-MSC), and cultured at 37 °C in a humidified atmosphere of 5% CO_2_ in serum-free medium for mesenchymal stem cells (NC0103+NC0103.S, Yocon, Beijing, China) as previously reported^36^ with some modification. When the proliferating colonies had reached near-confluence, the hESC-MSCs were passaged using Stem cells moderate digestive enzymes (NC1004, Yocon, Beijing, China). After 3-4 passages, MSCs were used for transplantation or other investigations. The CD immunoprofile of the hESC-MSCs was analyzed using flow cytometry to make sure that the obtained cells have the MSC phenotypes (CD73^+^, CD90^+^, CD105^+^, CD14^-^, CD34^-^, CD45-, CD79a^-^, and HLA-DR^-^)^37^.

### Materials

Ellman’s reagent (DTNB), cysteine, L-cysteine, and sulfosuccinimidyl 4-(N-maleimidomethyl) cyclohexane-1-carboxylate (sulfo-SMCC) were purchased from Thermo Scientific (USA). Human BDNF, recombinant human vascular endothelial GF (VEGF), and recombinant human basic fibroblast GF (bFGF) were purchased from PeproTech (USA). The bFGF ELISA Kit was purchased from MultiSciences (Hangzhou, China). DMEM was purchased from Corning (USA), and the EGM-e Bullet Kit was purchased from Lonza (USA). The Cell Proliferation Kit I (MTT) was purchased from Roche (Switzerland). Chicken anti-GFAP [abcam (ab134436), 1:500], rabbit anti-neurofilament (NF) heavy polypeptide [abcam (ab8135), 1:500], goat anti 5-HT antibody [Invitrogen(pa1-36157), 1:500], rabbit anti-MBP [ab218011], goat antiIBA1 [abcam (ab5076), 1:500], rabbit anti-NeuN [abcam (ab177487), 1:500], rabbit anti-fibronectin (FN) [abcam (ab23751, 1:500), and mouse anti-CD68 [abcam (ab201340, 1:500] were used. Goat anti-chicken IgY (H+L) secondary antibody conjugated with Alexa Fluor 488, donkey anti-rabbit IgG (H+L) highly cross-adsorbed secondary antibody conjugated with Alexa Fluor 555, donkey anti-mouse secondary antibody conjugated with Alexa Fluor, donkey anti-goat secondary antibody conjugated with FITC 488, and rabbit anti-goat secondary antibody conjugated with CY3 were purchased from Abcam (USA). Adenovirus-associated virus, AAV2/9-mCherry, was generated at the viral core of Zhejiang University, and its titer was adjusted to 1 × 10^13^ copies per mL before injection. Type A gelatin from porcine skin (300 bloom), 1-ethyl-3-[3-dimethylaminopropyl] carbodiimide hydrochloride (EDC), ethylenediamine (≥99.0%), 2-iminothiolane (≥98.0%), methacrylic anhydride (≥94%), triethanolamine (TEA, ≥ 99.0%), N-vinylcaprolactam (VC, ≥ 98%), Eosin Y (~99%), 5,50-dithiobis(2-nitrobenzoic acid) (DTNB) (≥98.0%), L-Cysteine (≥97.0%) and sodium borohydride (≥ 98.0%) were purchased from Sigma-Aldrich (USA). 1- (2-Aminoethyl)-1H-pyrrole-2,5-dione hydrochloride (NH2-MAL) was purchased from Aikonchemistry (Jiangsu, China). MPSS was purchased from Aladdin (Shanghai, China). Acrylate-poly(ethylene glycol)-N-hydroxysuccinimidyl-ester (ACLT-PEG-NHS, 3.5 kDa), 2- arm poly(ethylene glycol)-maleimide (2a-PEG-MAL, 20 kDa) was obtained from Tansh-Tech (Guangzhou, China).

### Synthesis and Characterization of Gelatin-SH

According to a reported procedure with minor modification^38^, gelatin type A was firstly aminated. In brief, one gram of gelatin was dissolved in 25 ml of 25 mL deionized (DI) water, and the pH was adjusted to about 5.1 using 1 M hydrochloric acid (HCl). After stirring at 40 °C for about 1 hour to confirm that gelatin was completely dissolved, ethylenediamine (3.14 mL) and a small amount of 1 M HCl were added to adjust pH to 5.0. And then, 0.5 g EDC was immediately added, and the mixture was stirred slowly at room temperature for 24 h. Then the mixture was transferred to a dialysis bag (MWCO 8-14 kDa) and dialyzed in deionized water in a dark room for 72 hours to remove impurities. The product was freeze-dried by a freeze-dryer to obtain a white floccular product, Gelatin-NH2, which was stored at 4 °C. Some products were taken for FT-IR and 1H-NMR characterization.

Next, for thiolation of the gelatin, a solution of Gel-NH2 (0.4 g) in DI water (20 mL) was prepared and adjusted to a pH of 7.0. Then add a slight excess of 2-iminothiacyclopentane hydrochloride and stir the reaction slowly away from light for 4 h. Afterwards, the mixture was transferred to a dialysis bag (MWCO 8-14 kDa) and dialyzed in 5 mM hydrochloric acid solution for 24 hours, in 1 mM hydrochloric acid solution for another 24 hours, and in DI water for a final 24 hours to remove impurities. A freeze-dryer subsequently freeze-dried the product to obtain a white flock-like product, some of which was taken for FT-IR and 1H-NMR characterization. The remaining product was stored at 4 °C for further investigation.

### Synthesis and Characterization of the Injectable Hydrogel

For formation of the injectable hydrogel, different concentrations of 2a-PEG-MAL (10%, 8%, 6%, 4% and 2%, wt%) was added to Gelatin-SH (8%, wt%). The gelling time was detected every 5 seconds. The elastic modulus was calculated from the linear portion of the stress-strain curve. Briefly, hydrogels with a uniform thickness of 1 cm were prepared by mixing 2a-PEG-MAL and Gelatin-SH of the determined concentration. The stress-strain curve was measured by a universal material testing machine (Roell Z020, Zwick, Germany) under a 50-N static load cell and a strain rate of 0.5 mm min^-1^. The degradation and swelling ratio of the hydrogel were evaluated using the following methods. Individual 2-mL hydrogel discs (5%) were incubated in 5 mL of PBS at 37 °C for 30 days to assess hydrolytic degradation. The mass of wet (Ms) and lyophilized (Md) gels were measured in triplicate at 0, 0.5, 1,3, 5, 7, 15, 30, and 60 days. To remove residual salt from the surface, the hydrogel was washed in distilled water before lyophilization. The total dry polymer mass loss of each sample was determined through comparison with the dry weight of the samples at 0 days. The swelling ratio was calculated as (Ms-Md)/Md at 0, 0.5, 1,3, 5, and 7 days.

### Synthesis of Maleimide-Modified, MP-Loaded liposomes (MP-lipo-MAL)

The MP-lipo-MAL was prepared using the thin film hydration method^39^. Firstly, (2, 3-dioleoxypropyl) trimethylammonium chloride (DOTAP), cholesterol, DSPE-PEG (2K) -Maleimide, DSPE-PEG, were all prepared into 5 mg/mL using chloroform. Then 840 μL DOTAP, 120 μL cholesterol, 50 μL DSPE-PEG (2K) –maleimide and 50 μL DSPE-PEG were added into a 50 mL pear-shaped bottle and mixed thoroughly. Afterwards, 1 mL of 4.2 mg/mL methylprednisolone dissolved in chloroform was added to the mixture. After the components are fully mixed, the mixture is rotary evaporated at 45 °C and −0.1MPa for 1-2 hours to form a uniform film and fully remove the chloroform. Following 1 mL of deionized water was added to the film, and ultrasonic hydration was performed for 30 min until the film completely fell off and a milky solution was obtained. Then, a lipid Extruder (Avanti^®^ Mini-Extruder) was used to push the ultrasonic hydration solution through 0.4 μm, 0.2 μm, and 0.1 μm filter membranes for 13-17 times at each level to obtain MP-lipo-MAL suspension. After lyophilization, the resulted MP-lipo-MAL was stored at 4 °C for further research.

### Measurement of the Physical Properties of MP-lipo-MAL

The MP-lipo-MAL’s particle size and zeta potential was determined using the dynamic light scattering technique (DLS, Zetasizer, Nano ZS, Malvern Instruments, UK). The shape and morphology of the MP-lipo-MAL were observed by scanning electron microscopy (SEM, Nova nano 450, Thermo, USA). Maleimide modification of the NPs was confirmed using Ellman’s test. MM-NPs (0.1 g) was added to 10 mL of L-cysteine (2 mM) and reacted for 2 h at RT. Then, the supernatant was collected by centrifugation (30 000 rpm, 60 min), and 4 μL of DTNB was added. After 30 min, the absorbance was measured by a microplate reader (Varioskan Flash, Thermo, USA), and the concentration of maleimide was calculated using a standard curve from 0–5 mM.

### Preparation of Injectable Hydrogel Solution

To prepare solution A, Gelatin-SH was dissolved in PBS (8%, wt%). To prepare solution B, 20% MP-lipo-MAL, maleimide-modified VEGF (10 ng/μL), -BDNF (50 ng/μL), and -bFGF (10 ng/μL) were added to 2a-PEG-MAL (5%, wt%) dissolved in PBS. Subsequently, solution B was mixed with solution A at a 1:1 ratio, resulting in MP-lipo-Gelatin.

## Evaluation of Cytotoxicity and Biocompatibility

### RNA isolation and qRT-PCR

Total RNA was isolated from cell lines and tissues using the TRIzol reagent according to the manufacturer’s instructions. cDNA was synthesized by using a reverse transcription kit (TAKARA, Japan) for mRNA. Quantitative expression was quantified using 2× SYBR Green qPCR Master Mix (Bimake, USA). The relative expression of mRNA was calculated with the method of 2-ΔΔCt and normalized by GAPDH. Each experiment was in triplicate^40^. Primer sequences are shown in Table S1.

### Sustained Release of MP and GFs

The release kinetics of MP and GFs from the hydrogel was studied as follows. One hundred microliters of MPG-HD were immersed in 5 mL of PBS and incubated at 37 °C. After 0, 0.5, 1, 3, 5, 7, 15, 30, and 60 days, the solution above the hydrogels was extracted and stored in Eppendorf tubes at −80 °C. The MP concentration was analyzed at 0, 1, 3, 5, and 7 days using high-performance liquid chromatography (LC-20, Shimazu, Japan) based on an existing procedure^41^. The bFGF concentration in the solution was measured using a bFGF ELISA kit according to the manual at 0, 0.5, 1,3, 5, 7, 15, 30, and 60 days. The absorbance at a wavelength of 450 nm was determined using a microplate reader.

### Surgical Procedure

Spinal contusion injury was induced using an Infinite Vertical impactor (68099, RWD Life Science, China). After the animals were anaesthetized with propofol (1 mL/100 g, injected intraperitoneally), laminectomy at the tenth thoracic vertebral level (T10-11) was performed by a dorsal laminectomy to expose the dorsal surface to induce spinal cord injury. Then, contusion was performed with a cylinder (diameter, 3 mm) that impacted the spinal cord at a rate of 2.5 m s^-1^ and stayed for 5 s. Artificial micturition was provided twice daily until the automatic restoration of bladder voiding. The following experiments were performed: 1) to evaluate the efficacy of intrathecal injection of MSCs in SCI, PBS or MSCs were injected intrathecally into SCI rats; 2) to test the efficacy of MP-embedded liposome and MSCs-loaded Hydrogel in SCI, PBS, MP-lipo-Gel, or MP-lipo-Gel + MSCs were injected into SCI rats.

Hydrogel (MP-lipo-Gel, and MP-lipo-Gel+MSCs) or PBS injection was performed 1 week after injury. A pulled-glass micropipette (68606, RWD, China) tipped 10-μL Hamilton microsyringe was used. The dorsal surface of the spinal cord with the contusive injury was re-exposed. After the injured area was identified, the hydrogel solutions with solution A were loaded in the micropipette and injected at the rate of 200 nl min^-1^ at 5 sites (1 μL per site). Two injections were performed at 2 different depths (0.6 and 1.2 mm). Solution B was injected at the same sites and under the same conditions. All injections were performed under the same conditions. AAV2/9-hsyn-mCherry injection for the anterograde tracing of propriospinal axons was performed at the T7-8 spinal cord, and AAV2/9-hsyn-mCherry was injected into the spinal cord at 80 nl min^-1^ using the following coordinates (150 nl per site): 0.4 and 0.8 mm lateral to the midline. Three injection sites were separated by 1 mm. At each injection site, 3 injections were performed at 3 different depths (0.5, 1.0, and 1.6 mm). The needle was left in every place for 1 min. Transection was performed at the injury site 9 weeks after the injury. Animals were sacrificed 8 or 10 weeks after injury.

### Histochemistry

Animals were sacrificed at 8 weeks after injury for histological assessment. Paraformaldehyde fixation was performed as previously described^34^. Twenty-micrometre-thick sections of the spinal cord were cut transversely using a cryostat (CryoStar NX50; Thermo, USA) and thaw-mounted onto Super Frost Plus slides (Fisher Scientific, USA). For immunohistochemistry, spinal cord tissue sections were incubated with primary antibodies overnight at 4 °C. After washing with PBS three times, the slides were incubated with appropriate secondary antibodies conjugated to fluorescent dyes. A fluorescence microscope (VS120, Olympus, Japan) was used to acquire images. The acquired images were further analysed using Image J software to evaluate the gray value of the indicated antibody and the cavity areas.

### Behavioral Assessment

The BBB scale assessed behavioral recovery for 10 weeks after injury in an open-field environment. Uninjured and sham-operated rats achieved full scores. Quantification was performed in a blinded manner. Detailed hindlimb kinematic analyses were performed based on data collected with the Vicon system. The method used for behavior recording was similar to a previously reported method^42^. Briefly, rats from different treatment groups were high lightened by anatomical landmarks on the hindlimb at the iliac crest, hip joint, knee joint, ankle and toe. Then, the animal was placed on a clear runway, and 8–10 continuous gait cycles were captured using the Vicon high-speed capture system for each rat. The captured videos were analyzed offline with Vicon Nexus software (Vicon Nexus motion system, UK). The iliac crest height and stride length were measured using landmarks, and stick views of hindlimb movement were drawn in MATLAB.

### Statistical Analysis

Data are shown as mean ± SEM or mean ± SD as indicated in the figure legends. Statistical significance of differences between two experimental groups was calculated using unpaired, two-tailed Student’s t test in GraphPad Prism 6.0 software and a *P*-value less than 0.05 was considered significant.

## Acknowledgments

This study was supported by the National Natural Science Foundation of China (81971866 to X.W.), the Scientific and Technological Innovation 2030 Program of China - major projects (2021ZD0200408 to X.W.), the Science Fund for Distinguished Young Scholars of Zhejiang Province (LR20H090002 to X.W.), the Leading Innovative and Entrepreneur Team Introduction Program of Zhejiang (2019R01007 to X.W.), the Fundamental Research Funds for the Central Universities (K20210195 to X.W.) and the Postdoctoral General Foundation project (2020M671746 to X.C.).

## Author Contributions

X.C., W.L., J.Y. and Y.Z. contributed equally to this work. X.W., and X.C. conceptualized and designed the study. X.C., W.L., J. Y., Z. W. and Y.Z. conducted the experiments and collected the data. X.C., J.Y., S.J., W.C., X.L., Z.W., B.Y., X.G. and X.W. analyzed and interpreted the data. X.C. and X.W. drafted the paper. All authors critically revised the manuscript and approved the final version for submission.

## Data Availability Statement

The data to support the findings of this study are included in the paper and supplementary information, and further data are available from the corresponding author upon reasonable request.

