## Supplemental Tables and Figures for "Injectable hydrogel embedded with mesenchymal stem cells repairs severe spinal cord injury"

1. Department of Neurobiology and Department of Orthopedics, 2nd Affiliated Hospital, Zhejiang University School of Medicine
2. NHC and CAMS Key Laboratory of Medical Neurobiology, MOE Frontier Science Center for Brain Research and Brain–Machine Integration, School of Brain Science and Brain Medicine, Zhejiang University, Hangzhou, Zhejiang Province, 310003, PR China
3. Department of Rehabilitation Medicine, First Affiliated Hospital, College of Medicine, Zhejiang University, Hangzhou, Zhejiang Province, 310003, PR China
4. Key Laboratory of Neuroregeneration of Jiangsu and Ministry of Education, NMPA Key Laboratory for Research and Evaluation of Tissue Engineering Technology Products, Nantong University, Nantong, China
5. Co-innovation Center of Neuroregeneration, Nantong University, Nantong, 226001 Jiangsu, PR China
6. These authors contributed equally
7. Lead contact

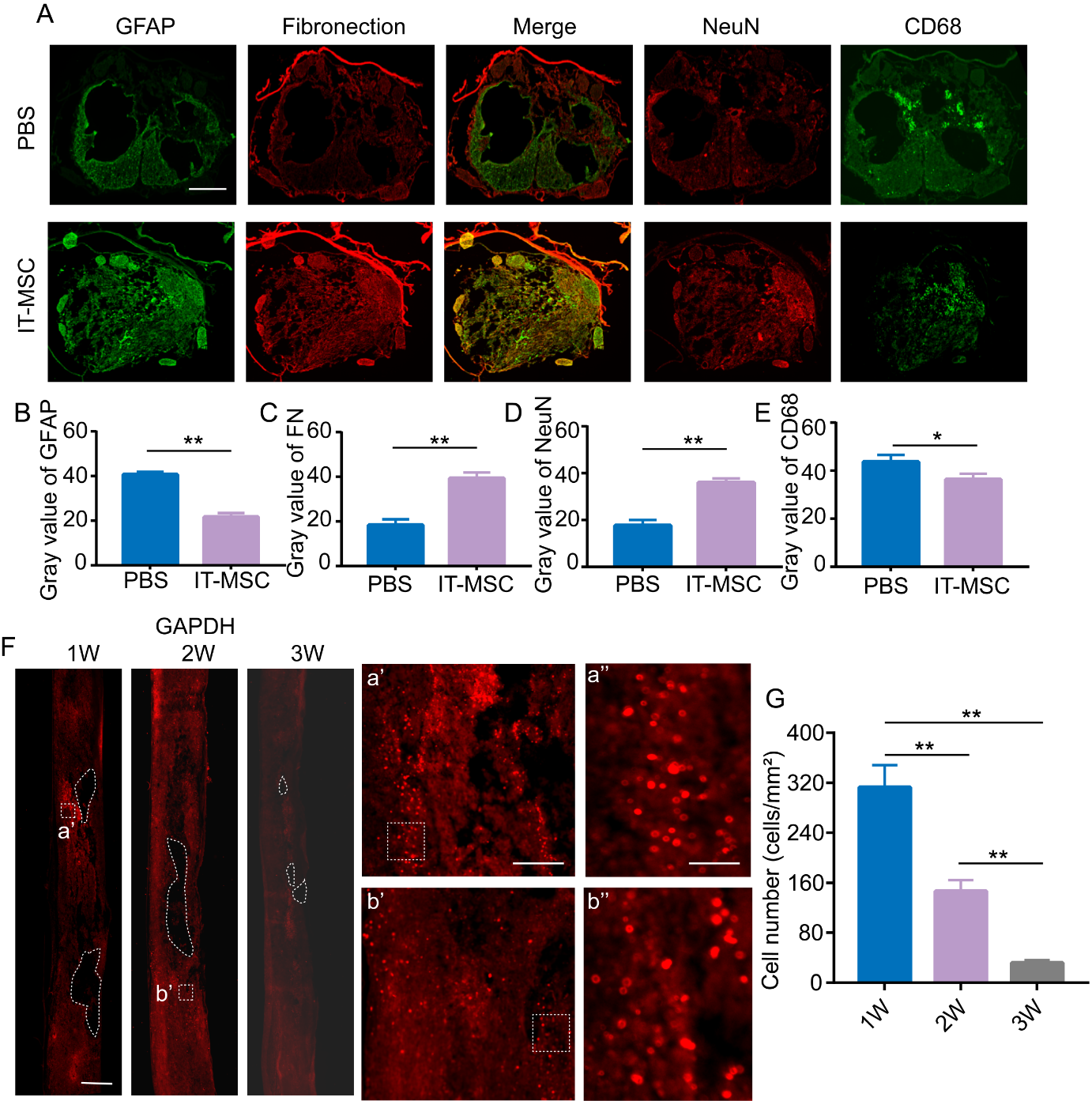

**Figure 1S. MSCs treatment through** **intrathecal injection reduced the inflammation effect at the injured site.**

(A) Representative images of immunofluorescent staining of Coronal sections at the injured sites for GFAP (green), Fibronection (red), NeuN (red) and CD68 (green) under different injection conditions after 8 weeks. Bar=100 μm.

(B-E) Quantification of GFAP (B), Fibronection (C), NeuN (D) and CD68 (E) density by the gray value; Data are presented as mean ± SD (n=5). Statistical analysis was performed using unpaired, two-tailed Student’s t test one-way ANOVA followed by Bonferroni's post hoc test. **P* < 0.05 and ***P* < 0.01.

(F) Representative images of immunofluorescent staining of Sagittal sections at the injured sites for human specific GAPDH (red) after 1, 2, 3 Weeks following intrathecal injection of MSCs. a’, b’, a” and b” show the details for MSCs survival at the injury site. Bar=100 μm.

(G) Quantification of survival cell numbers at the injury sites by counting; Data are presented as mean ± SD (n=3). Statistical analysis was performed using unpaired, two-tailed Student’s t test one-way ANOVA followed by Bonferroni's post hoc test. ***P* < 0.01.

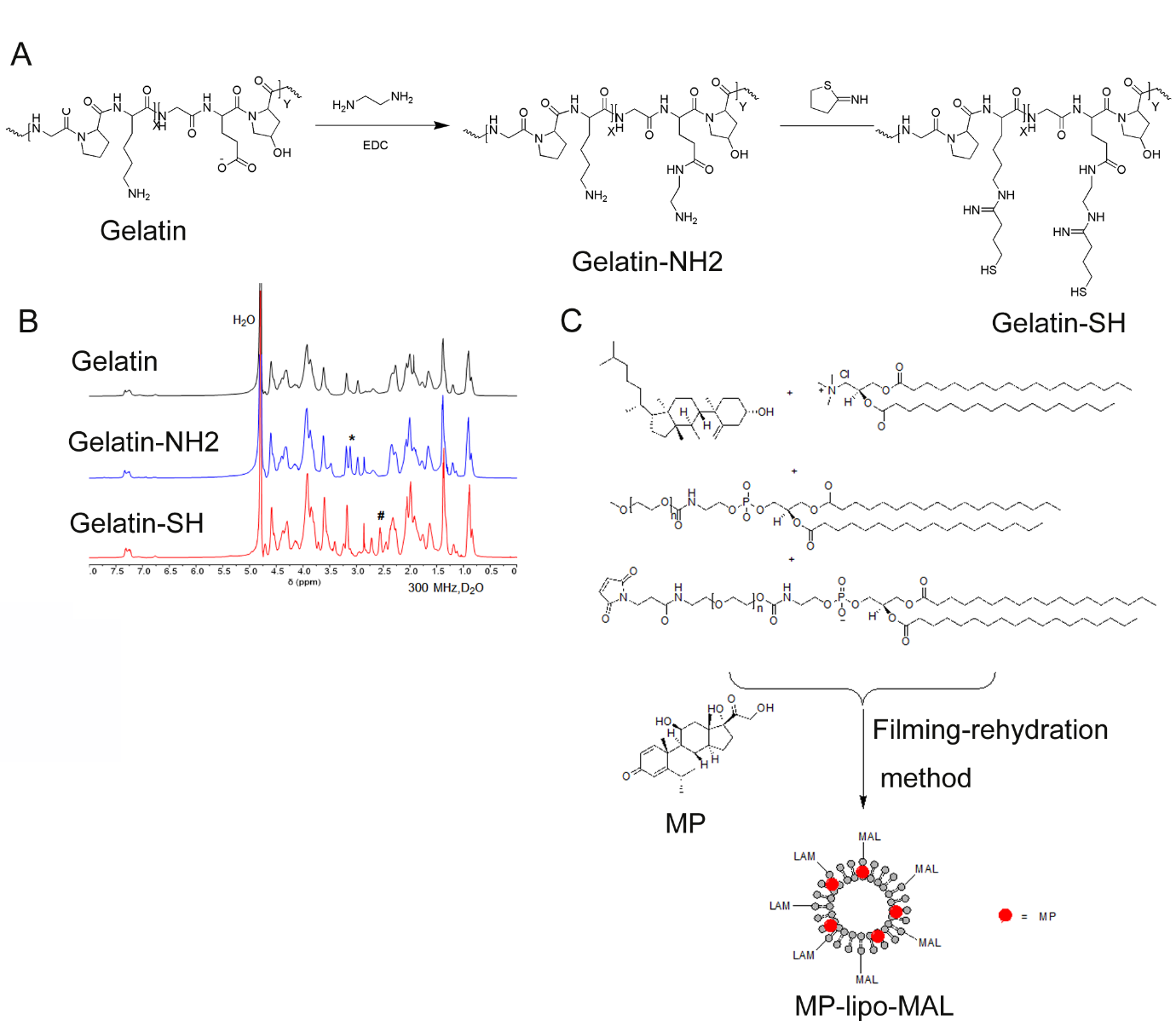

**Figure 2S. Synthesis of related materials and nanoparticles.**

(A) Synthesis equation of Mercaptoization for gelatin.

(B) The NMR maps of Gelatin, Gelatin-NH2 and Gelatin-SH.

(C) Synthesis flowchart of Maleimide-Modified, MP-Loaded liposomes (MP-lipo-MAL).

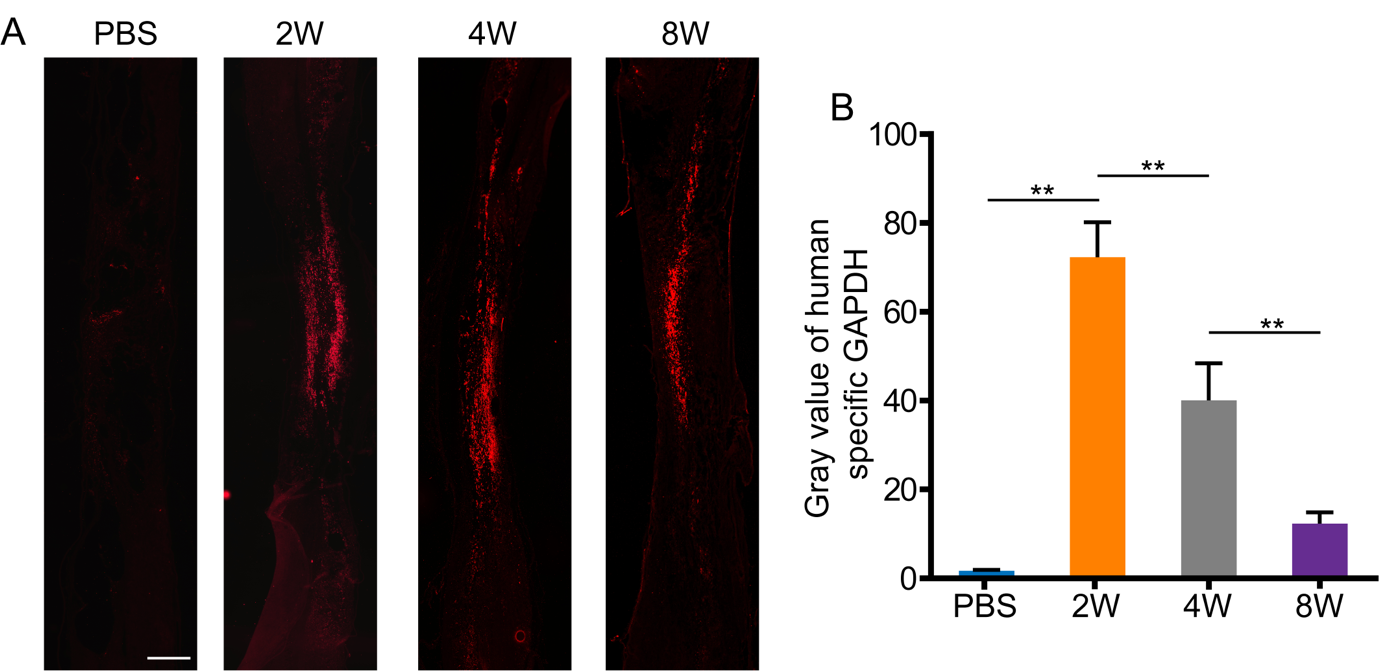

**Figure 3S. The survival analysis of MSCs at the injury sites after in situ injection.**

(A) Representative images of immunofluorescent staining of Sagittal sections at the injured sites for human specific GAPDH (red) after 2, 4 and 8 Weeks following in situ injection of Gel+MSCs. Bar=100 μm.

(B) Quantification of survival cells at the injury sites by the gray value; data are presented as mean ± SD (n=3). Statistical analysis was performed using unpaired, two-tailed Student’s t test one-way ANOVA followed by Bonferroni's post hoc test. ***P* < 0.01.

**Table S1. Primer sequences used for real-time qPCR.**

| Gene name | Forward (5’-3’) | Reverse (5’-3’) |
| --- | --- | --- |
| GAPDH（human） | ACCCAGAAGACTGTGGATGG | TCAGCTCAGGGATGACCTTG |
| SOX2（human） | AATGCCTTCATGGTGTGGTC | GCTTAGCCTCGTCGATGAAC |
| OCT4（human） | CGAGTGTGGTTCTGTAACCG | CTGAGAAAGGAGACCCAGCA |
| NANO（human） | GCTCTTACTGACTGGCATGAG | CGCAGCTCTAGGAGCATGTG |
| GAPDH（rat） | CAAGGCTGAGAATGGGAAGC | GAAGACGCCAGTAGACTCCA |
| IL-6（rat） | CCACTGCCTTCCCTACTTCA | ACAGTGCATCATCGCTGTTC |
| TNF-α（rat） | GGTCCCAACAAGGAGGAGAA | GCTTGGTGGTTTGCTACGAC |
| IL-4（rat） | CTTGCTGTCACCCTGTTCTG | ACAAACATCTCGGTGCATGG |
| IL-13（rat） | CCCTCAGGGAGCTTATCGAG | ATACCATGCTGCTGTTGCAC |
